# Macroscale Cortical Organization and a Default-Like Transmodal Apex Network in the Marmoset Monkey

**DOI:** 10.1101/415141

**Authors:** Randy L. Buckner, Daniel S. Margulies

## Abstract

Networks of widely distributed regions populate human association cortex. One network, often called the default network, is positioned at the apex of a gradient of sequential networks that radiate outward from primary cortex. Here extensive anatomical data made available through the Marmoset Brain Architecture Project were explored to determine if a homologue exists in marmoset. Results revealed that a gradient of networks extend outward from primary cortex to progressively higher-order transmodal association cortex in both frontal and temporal cortex. The transmodal apex network comprises frontopolar and rostral temporal association cortex, parahippocampal areas TH / TF, the ventral posterior midline, and lateral parietal association cortex. The positioning of this network in the gradient and its composition of areas make it a candidate homologue to the human default network. That the marmoset, a physiologically- and genetically-accessible primate, might possess a default-network-like candidate creates opportunities for study of higher cognitive and social functions.

---

The common marmoset, *Callithrix jacchus*, is a small New World primate that is increasingly being chosen as a model system for neuropsychiatric illness and studies of higher cognitive and social functions (Belmonte et al., 2015; Miller et al., 2016; Jennings et al., 2016; Okano et al., 2016; Oikonomidis et al., 2017; Silva, 2017). One reason to turn to a primate to complement other available rodent models is that primates have a clear prefrontal cortex including granular and dysgranular regions (Öngür & Price, 2000; Wise, 2008; see Preuss, 1995 for discussion). For example, histological and connectional studies of frontopolar cortex note that marmosets (Burman et al., 2006; Burman & Rosa, 2009) possess a granular area 10 sharing properties with the large-brained New World (Rosa et al., 2018) and Old World (Carmichael and Price, 1994; Petrides & Pandya, 2007) monkeys as well as the human (Petrides & Pandya, 1994; Öngür et al., 2003). Area 10 falls at the rostral apex of prefrontal cortex and is implicated in advanced forms of planning, abstract reasoning, and handling multiple completing task demands (Ramnani and Owen, 2004; Tsujimoto, Genovesio, & Wise, 2011; Mansouri et al., 2017). Area 10, particularly the medial extent of area 10, is a consistent node in the human default network (Buckner et al., 2008), raising the interesting possibility that conserved network organization in the marmoset may include a homologue of the default network.

However, the marmoset is a particularly small primate with a brain 180th the size of the human brain. It’s frontopolar region is comprised of a relatively homogenous area 10 with elements of a gradient rather than the more clearly differentiated architectonic subfields observed in the macaque and human (Carmichael & Price, 1994; Petrides & Pandya, 1994; Öngür et al., 2003; Burman & Rosa, 2009). Thus, the marmoset has intriguing relations to the human but its evolutionary distance (41-46 million years since the last common ancestor with human; Perelman et al., 2011; Hedges et al., 2015), small brain size, and less differentiated prefrontal cortex raise uncertainty about how much homology should be expected. Two major features of large-scale network organization raise questions about homology that can be explored through analysis of anatomical connectivity.

Human association cortex is populated by a series of large-scale networks (Yeo et al., 2011; Power et al., 2011). Multiple separable networks include canonical sensory-motor networks through to the widely distributed association networks. In terms of topology, the multiple networks form an orderly progression that radiates outwards from sensory cortex to transmodal association cortex (Margulies et al., 2016; Braga & Buckner, 2017; Huntenberg et al., 2018). *A first open question is whether the marmoset possesses a macroscale gradient among networks similar to the human.*

Situated at the farthest end of the macroscale gradient of networks in the human is the default network (Margulies et al., 2016; Braga & Buckner, 2017). The default network behaves in peculiar ways as compared to other well-studied cortical networks. In particular, the default network increases activity when attention to the external environment is relaxed and internal, constructive modes of cognition emerge (see Gusnard & Raichle, 2001; Buckner et al., 2008; Andrews-Hanna et al., 2014 for reviews). It is also active during directed tasks involving remembering and social inferences drawing a great deal of interest (Spreng et al., 2009). *For these reasons, a second open question is whether the marmoset possesses a default-network-like candidate.*

The human default network comprises widely distributed regions including (I) medial prefrontal cortex extending from the frontal pole to the anterior cingulate, (II) precuneus, posterior cingulate and retrosplenial cortex, (III) a caudal region of the inferior parietal lobule, (IV) temporal association cortex extending into the temporal pole, and (V) parahippocampal cortex (Greicius et al., 2003; Buckner et al., 2008; Raichle, 2015). Recent analyses of individual brains reveal further network subdivisions and topographic details. The canonical default network is most likely multiple interwoven networks and contains additional regions beyond the five mentioned above, including dorsolateral and ventrolateral prefrontal cortex (Braga & Buckner, 2017; see also Gordon et al., 2017; Kong et al., 2018). At a coarse scale, the five highlighted zones are repeatedly identified in analyses of human neuroimaging data and can serve as an anchor for identifying a candidate homologue, specifically the network recently labeled default network – A (Braga & Buckner, 2017). Default network – A is distinguished from the spatially adjacent default network – B by strong correlation with parahippocampal and retrosplenial cortex. Moreover, there is evidence in the macaque for homology suggesting the network was present in a last common ancestor ∼28-31 million years ago (Vincent et al., 2007; Buckner et al., 2008; Margulies et al., 2009; Binder et al., 2009; Ghahremani et al., 2017).

A candidate homologue of default network - A is hypothesized to have three anatomical properties. First, the candidate should include regions at the transmodal apex (Buckner & Krienen, 2013; Margulies et al., 2016; Braga & Buckner, 2017; Hunterberg et al., 2018). Anatomically, a gradient of networks is expected moving outward from caudal frontal cortex (primary motor cortex) through to frontopolar cortex. The default network is hypothesized to include areas situated at the rostral apex. Second, the candidate should comprise at least the five distributed zones of cortex discussed above and previously described in detail in relation to architectonic areas (Buckner et al., 2008; see also Margulies et al., 2009). Based on estimated architectonic correspondence, these five zones in the marmoset are predicted to include regions at or near: (I) frontopolar A10, (II) posterior midline A29a-c, A23, caudal A30, (III) rostral temporal association cortex TE3 / TPO / PGa/IPa, (IV) posterior parietal cortex Opt / PG, and (IV) parahippocampal cortex TH/ TF. Some of these fields are expanded and differentiated into subfields in macaque and human, so they may also be expected to encompass a relatively smaller portion of the marmoset brain (see Chaplin et al., 2013).

Third, the candidate should be anatomically distinct from canonical sensory-motor hierarchies, in particular the network involving FEF and the MT complex. The reason for this third expectation is that, in the human the two networks are functionally antagonistic (Fransson et al., 2006; Fox et al., 2005). Thus, while anatomical tract tracing studies have often interpreted higher-order association regions as polysensory convergence zones (e.g., Jones & Powell, 1970), network analysis in the human provides evidence for separate parallel networks (see also Goldman-Rakic, 1988; Mesulam, 1981; 1990 for discussion). In Petrides and Pandya’s (2007) analysis of the connectivity of the frontal pole in macaque, the absence of connectivity with regions participating in visuospatial and motor demands led to the conclusion that macaque area 10 “does not regulate attention to events occurring in external space”.

In the present paper, extensive tract tracing data from the Marmoset Brain Architecture Project were explored to determine (1) whether there is a macroscale gradient of multiple networks in the marmoset that progresses from sensory zones to an apex transmodal network and (2) whether the apex transmodal network has properties to suggest provisional homology with the human default network.

## Results

### A gradient of networks extends outward from somatomotor cortex to progressively higher-order transmodal association cortex

The sequential pairings of frontal and posterior cortical injections suggest multiple distinct potentially parallel networks (Figures 1 & 2). These multiple networks are spatially near to one another along a caudal to rostral sequence in frontal cortex.

**Figure 1.**
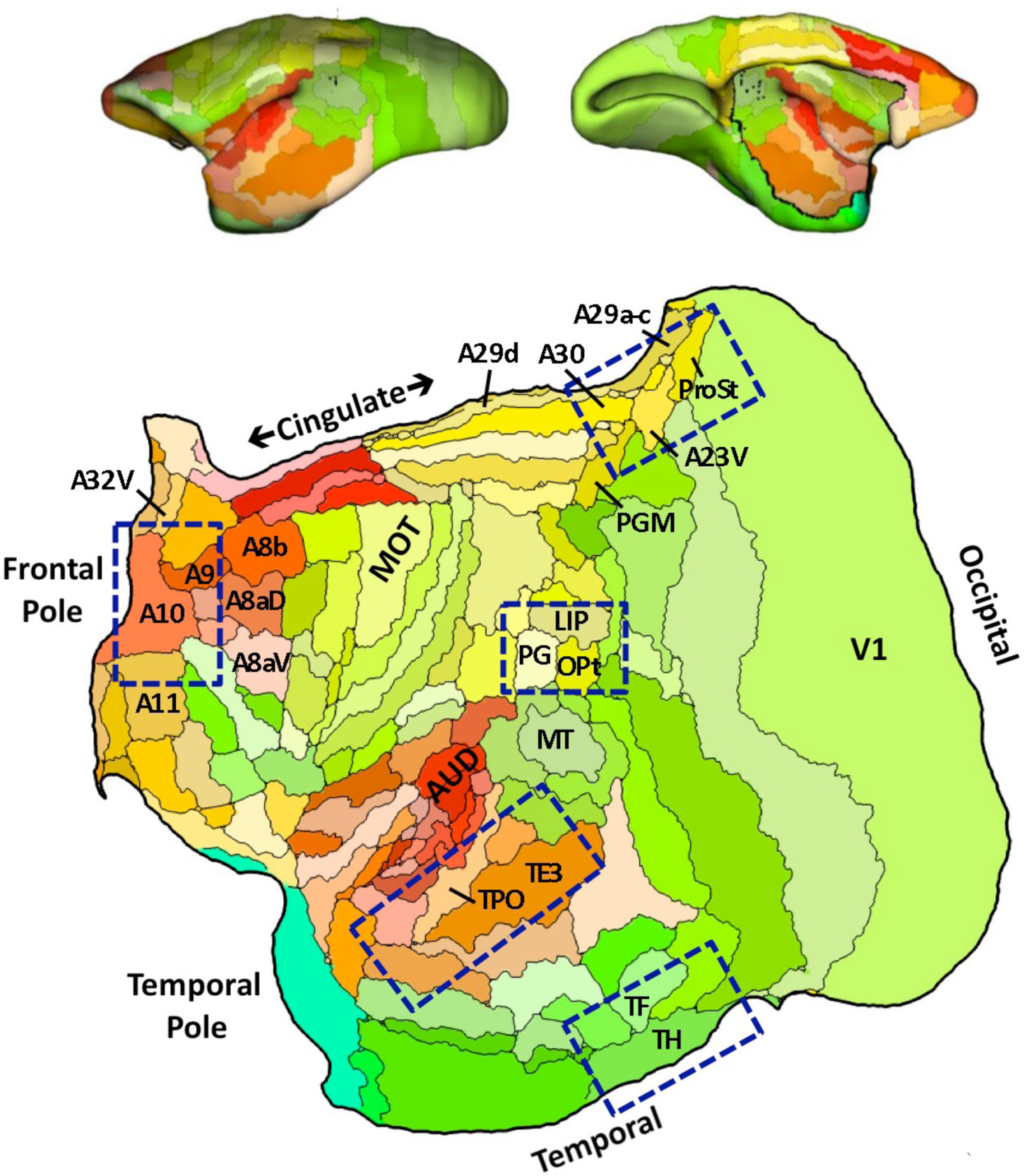
Flat map format. All cortical areas are displayed on a flat map that minimizes distortion (Majka et al., 2016). The lateral (top left) and midline (top right) show the volume surface models of the marmoset cortex color-coded corresponding to the flat map representation below. Relevant areas are labeled for orientation as well as MOT (primary motor cortex A4ab) and AUD (auditory cortex involving multiple primary auditory areas). The major zones of interest in this paper are highlighted by blue rectangles. These are not the only zones implicated in the default network but represent five zones that are candidate homologues to several of the well-studied regions implicated in default network – A in the human (Braga & Buckner, 2017). Area labels use nomenclature of the Paxinos et al. (2012) atlas.

**Figure 2.**
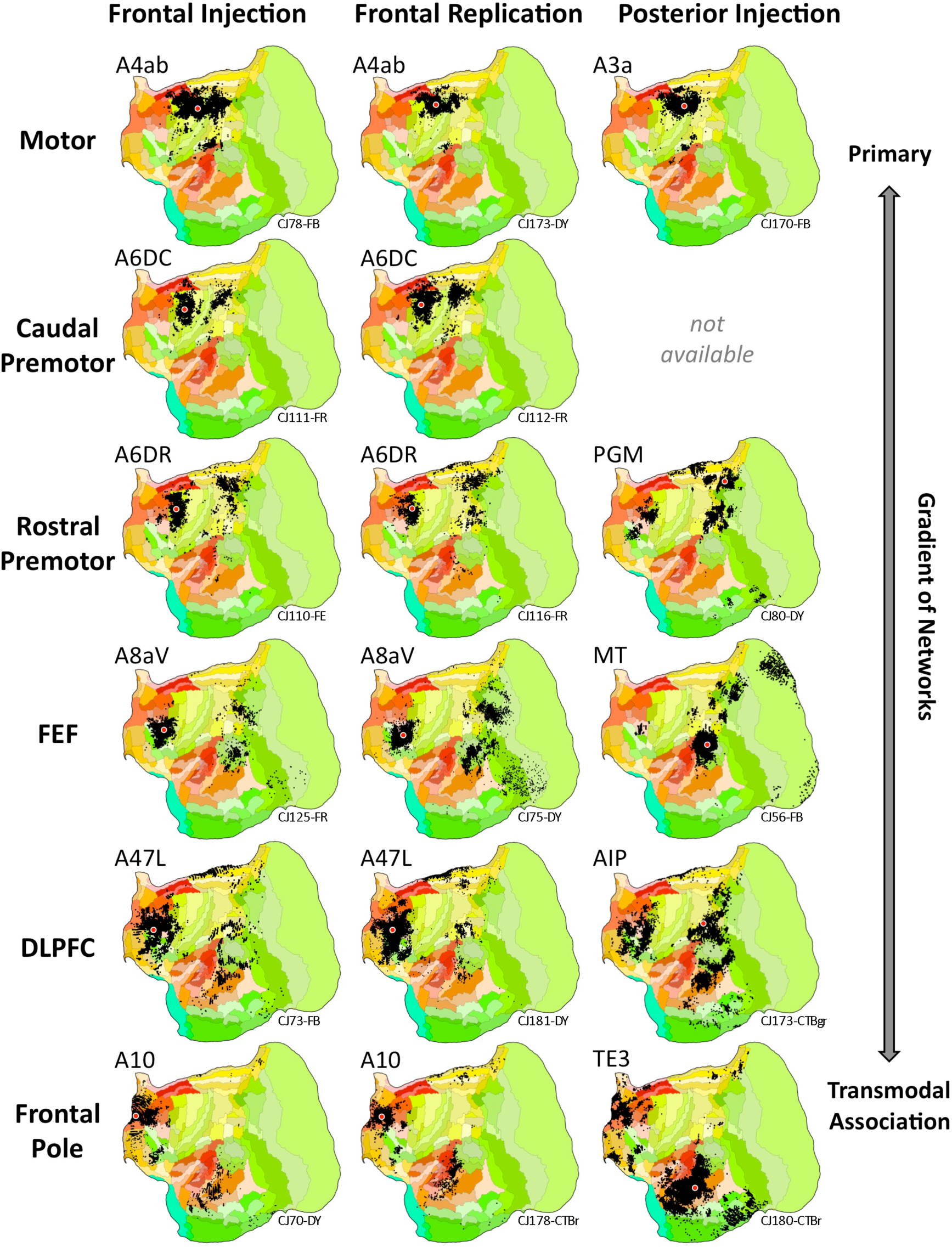
Aggregate analysis of anatomical connectivity reveals a macroscale gradient of networks. Each row displays a candidate network that is based on a frontal injection (left column), replication of the frontal injection (middle column), and corroboration of the network using a posterior injection (right column). Flat map format from Figure 1. These displayed networks represent a partial subset of possible networks that could be plotted and do not reflect the full complexity of the projection patterns. Nonetheless, they reveal a macroscale gradient by which progressively more distributed networks populate the cortex as one goes from primary motor cortex (A4ab) through to frontopolar A10. In each map, the tracer injection is shown with a red dot (all retrograde). Injection cases are labeled in the bottom right as annotated in the Marmoset Brain Architecture Project archive.

The networks beginning with A6DR-PGM are more widespread than the lower level somatomotor networks (A4ab-A3a and A6DC) including projections from midcingulate, temporal visual and transmodal association cortex. The complexity of this distributed pattern and that of the A8aV-MT network make them difficult to sequence relative to one another. They both possess features that position them lower in the macroscale gradient of networks relative to the A47L-AIP and A10-TE3 networks. Their connections to frontopolar cortex are minimal or absent and connections to rostral temporal association cortex are also restricted. Our ordering reflects that A8aV is rostral to A6DR.

As the sequence progresses, the network involving A47L and AIP prominently includes extensive regions of prefrontal, temporal and parietal association areas. The posterior projections to A47L spare MT and spread rostrally into temporal cortex. Similarly, the projections from posterior parietal cortex are broadly rostral to those involved in the A8aV – MT network, possibly capturing a feature observed in human homologues (e.g., Vincent et al., 2008).

The apex transmodal network involving A10 encompasses many zones expected of a default network candidate (see also Burman et al., 2011). A10 receives projections from extended regions of rostral temporal association cortex extending toward the temporal pole. Even when additional prefrontal injections were sought at or near A47L for contrast, none could be found that included comparable connections to the most rostral zones of temporal association cortex (e.g., see Case CJ800-CTBgr injection in A45 with some invasion of A47). The connections to the posterior injection (in rostral TE3) include A10 and surrounding regions, as well as prominent label in the parahippocampal region (TH / TF), modest projections from posterior cingulate / retrosplenial cortex and a zone in posterior parietal cortex on the border of Opt, PG, and LIP.

In addition to providing evidence for a sequence of progressively more distributed networks that situate themselves along a caudal to rostral frontal gradient, these composite patterns suggest a default-network-like candidate.

### The apex transmodal network is a candidate homologue of the human default network

The pattern of connections to A10 is consistent with it being a component of a default network - A homologue. Figure 3 illustrates the anchor A10 case from Figure 1 in greater detail as well as five additional cases to illustrate common patterns. The first observation is that the broad pattern is similar across several A10 injections and extends to nearby areas (A11, A9, and A32V), all receiving projections from rostral temporal association cortex, posterior cingulate / retrosplenial cortex, and parahippocampal cortex (TH and sometimes TF). The posterior cingulate projections extend into area prostriata (ProSt) and in some cases ProSt is more densely labeled in particular near to the border of A23V.

**Figure 3.**
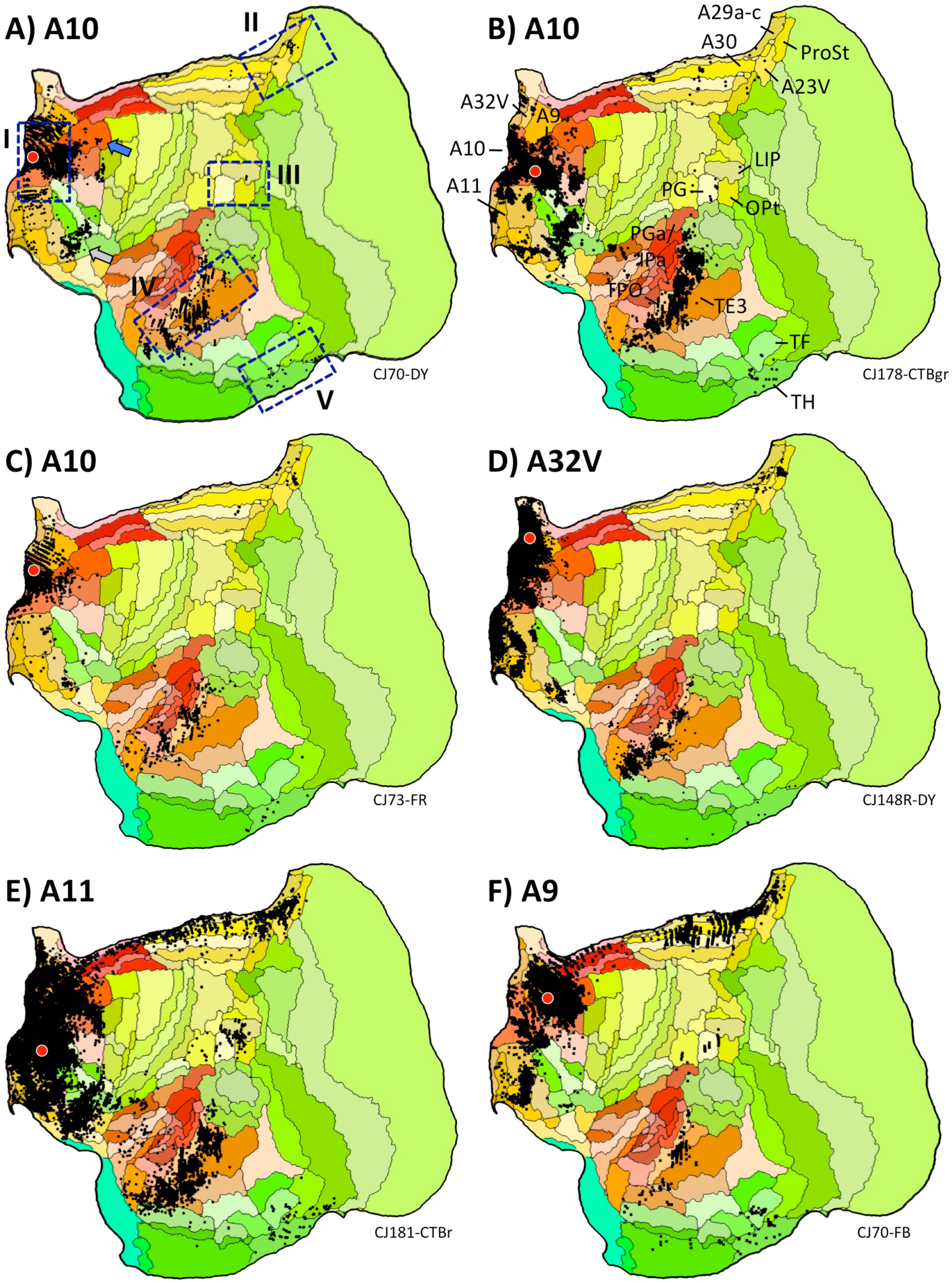
Anatomical evidence for a default-network-like candidate in the marmoset. Multiple injections from frontopolar cortex A10 and adjacent areas are illustrated. The blue bounding boxes in A identify five target zones hypothesized to be part of the default network – A, labeled I to V. The area labels in B are displayed for key areas hypothesized to be components of default network – A. The blue arrow in A notes projections originating in A8ab, which is examined in more detail in Figure 5. The gray arrow in A notes projections within and along the border of A47O/A13L/A13M that are discussed in the text. Arrows also illustrate similar patterns from temporal injections in Figure 4E.

While frontopolar A10 is generally considered to be absent of direct projections from posterior parietal cortex, weak projections are present in some cases from the region of Opt/PG/LIP. When present the labeling is quite modest (e.g., Cases CJ170-DY and CJ178-CTBgr). Injection to A11, the interconnected area adjacent to A10 on the orbital surface, and A9, the area adjacent on the dorsomedial surface, show clear projections from Opt/PG/LIP while also recapitulating much of the connectional pattern of A10 (Cases CJ181-CTBr and CJ70-FB).

The projections to temporal association cortex provide convergent evidence with the frontopolar injections for a default network homologue. An injection within rostral TE3 was the original target (Figure 2). Based on its pattern, adjacent TPO was explored more thoroughly. TPO, possibly related to macaque superior temporal polysensory area defined on physiological properties (Cusick et al., 1995), is near to auditory cortical areas (e.g., AuCPB and AuML). This complex region thus borders higher order auditory processing areas as well as transmodal candidates of the default network. Multiple lateral temporal lobe injections were examined that contrasted the ‘default-network-like” pattern of caudal TPO and rostral TE3 with nearby injections in auditory areas (AuCPB and AuML). The results are displayed in Figure 4.

**Figure 4.**
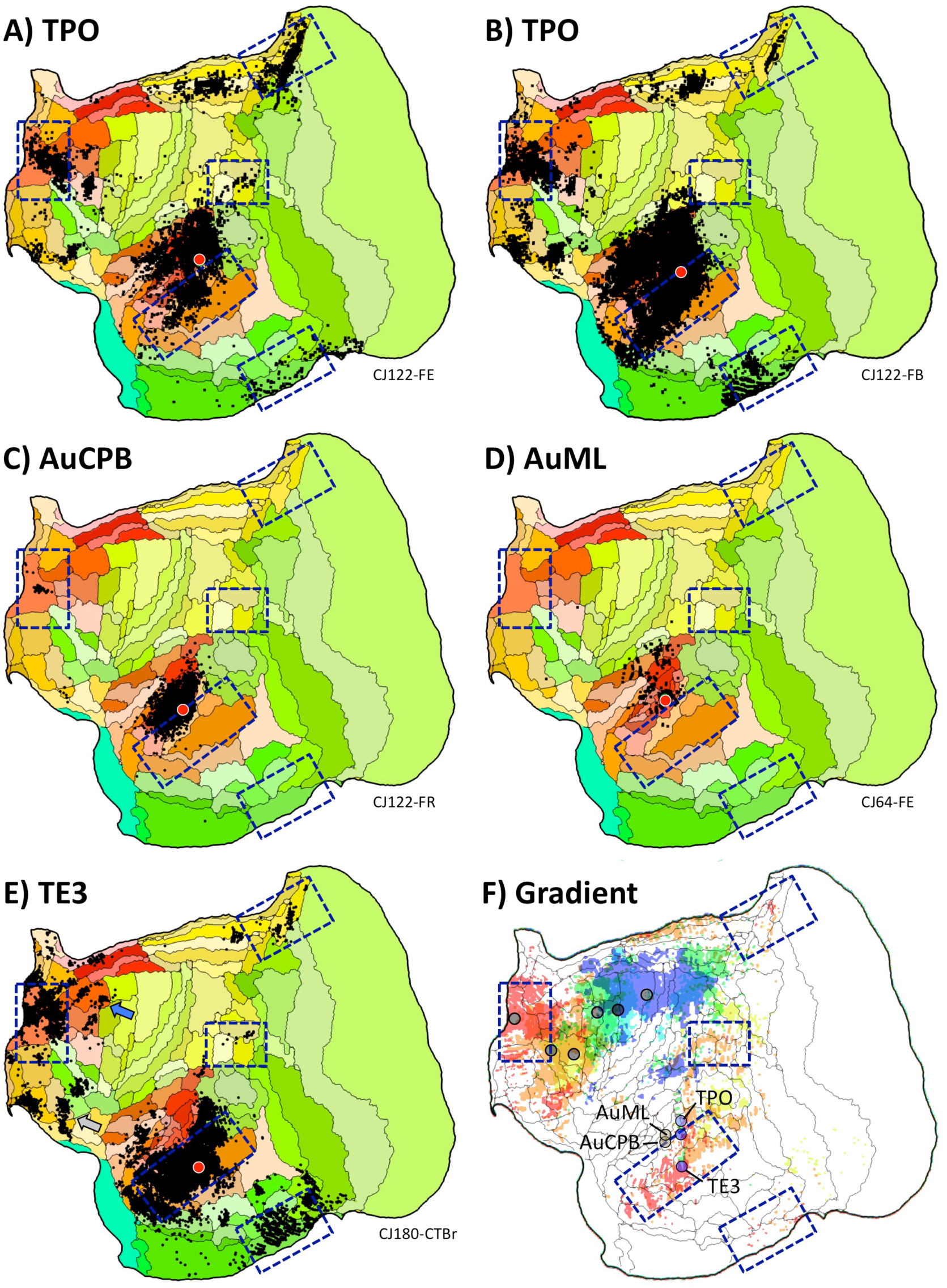
Evidence for involvement of temporal association cortex in the default-network-like candidate. Injections in estimated TPO (A, B) are distinct from injections to nearby auditory areas AuCPB and AuML (C, D). TPO injections receive projections from the full constellation of regions predicted as components of the default network – A candidate as well as nearby auditory areas, whereas the auditory areas receive projections predominantly from local auditory areas. (E) Injection of TE3 also yields a default-network-like pattern, while largely sparing auditory cortex. (F) The gradient of temporal lobe projections to frontal regions is illustrated by combining the projection patterns from the left column of Figure 2 into a single multicolored image: A4ab, purple; A6DC, blue; A6DR, green; A8aV, yellow; A47L, orange; A10, red. A clear progression into rostral temporal cortex is observed. The three blue circles mark the locations of the TPO and TE3 injections and the two tan circles mark the auditory cortex injections (AuCPB and AuML). In addition to revealing the network gradient, this composite image illustrates a region of temporal cortex where transmodal association areas that are part of the default-network-like candidate are near to auditory sensory areas (see also Supplementary Figure 2).

The TE3 injection recapitulates nearly the full extent of the candidate default network homologue (Case 180-CTBr). The caudal TPO injections (CJ122-FE and CJ122-FB) display a broad set of projections overlapping much of the zones involved in the candidate default network homologue as well as extensive local projections from auditory cortex. By contrast, the nearby injections in auditory areas AuCPB (Case CJ122-FR) and AuML (Case CJ64-FE) are predominantly local, avoiding nearly all of the default network zones. The one exception was involvement of A10 for CJ122-FR, but it should also be noted that injection, while primarily restricted to the caudal parabelt, may have involved the anterior lateral belt as well.

Figure 4F plots the locations of the temporal injections that yield the default-network-like projection pattern overlaid onto the projection patterns of the caudal-to-rostral frontal injections. The caudal TPO and rostral TE3 injections fall within or on the border of the zones linked to the apex transmodal network. The rostral TE3 injection is fully within the region of frontopolar projections, while the caudal TPO injections are on the border. Of further note is the global observation that the gradient of temporal lobe projections to frontal cortex does not respect the areal boundaries. The projection gradient goes through caudal TPO and rostral TE3, sparing large portions of the two areas.

### Evidence for separation of the apex transmodal network from a sensory-motor network

The distributed network associated with A10-TE3 is spatially separable from the A8aV-MT network across the brain (Figure 2). To illustrate this point, bounding boxes marking the five zones of the candidate default network homologue were placed on high-resolution views of injection patterns for frontal A8aV (Case CJ125-FR) and posterior MT (Case CJ56-FB)(Supplementary Figure 1). What is notable is that candidate default network zones are largely absent from participation in the A8aV-MT network.

### The apex transmodal network is embedded within a complex, distributed organization

The analyses have focused on finding a candidate marmoset homologue of the human default network anchoring on five defining zones. These zones are a subset of the complex network in the human, preferentially targeting components of what has recently been called default network – A (Braga & Buckner, 2017). That network shows strong coupling to the parahippocampal cortex and ventral portions of the posterior midline and retrospenial cortex. The marmoset candidate described to this point has these properties. Motivated by recent work of Liu et al. (submitted), additional tracer injections explored marmoset areas A8b (Cases CJ73-DY and CJ83-DY) and A8aD (CJ800-CTBr and CJ108-FR) in relation to the posterior midline area PGM. One injection already described in Figure 2 was within PGM fully (CJ80-DY). An additional injection bordered A23V (Case CJ84-FB).

Figure 5 shows the results. The patterns are notable in that projections to A8b and, to a lesser extent, A8aD overlap or are adjacent to zones highlighted by the analyses of the A10-TE3 network including projections from posterior cingulate, temporal association cortex near to TPO and PGa/IPa, parahippocampal cortex TH / TF, and a zone near to the Opt / PG / LIP cluster. As illustrated in Figure 3 the A10-TE3 network receives projections from A8b and A8aD (see also summary Figure 2 of Burman et al., 2011). The injections to PGM display less compelling involvement. In particular the injection fully in PGM has minimal projections to the posterior cingulate and retrosplenial regions implicated in the default network – A candidate. The PGM injection that shows the most integration is Case CJ84-FB, which falls on the border with A23V and A30.

**Figure 5.**
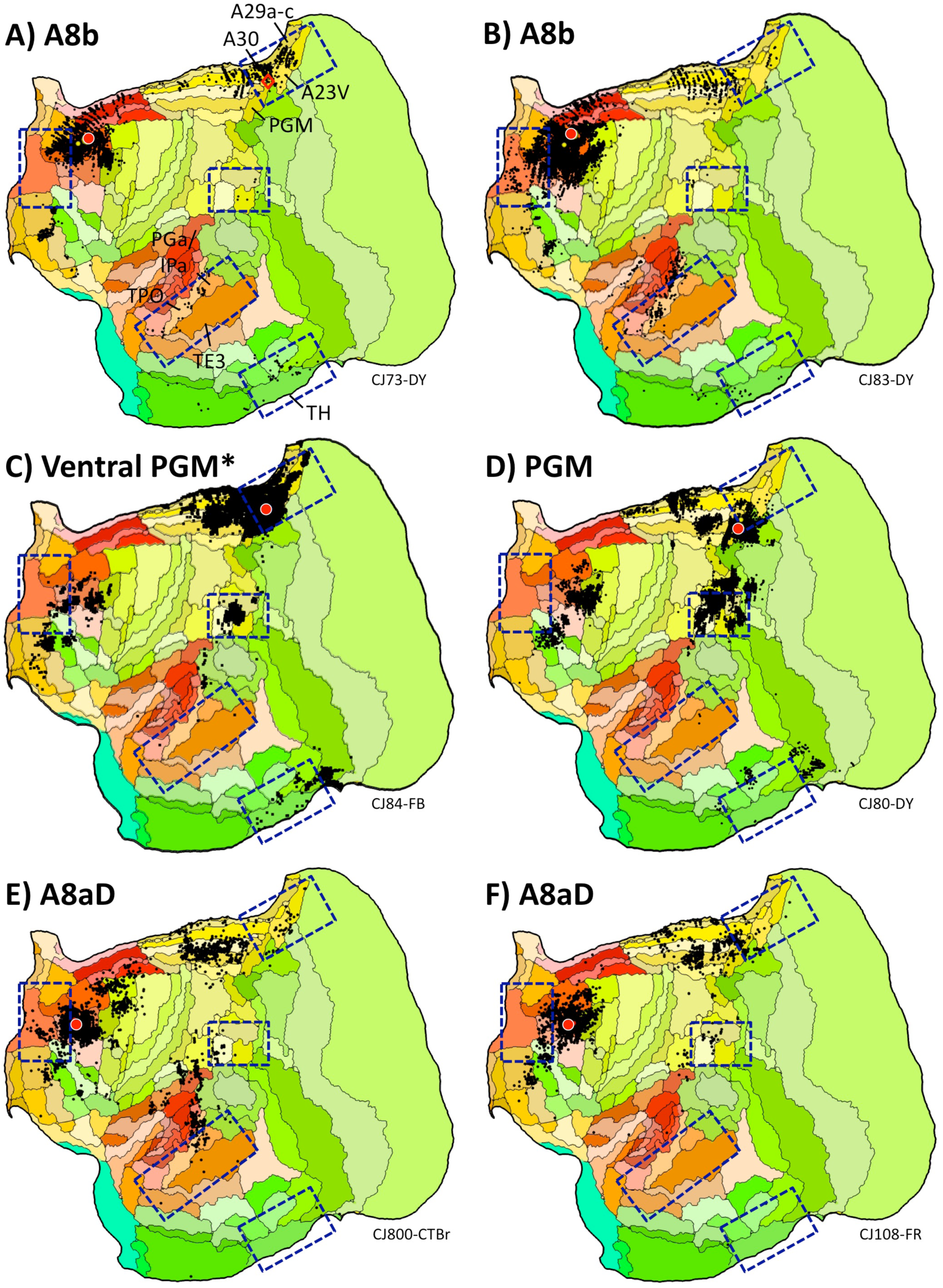
Additional areas may be components or subdivisions of the default-network-like candidate. The human default network contains regions outside of the five-targeted zones, including additional prefrontal regions and potentially more extensive areas along the posterior midline. Injections to two relevant frontal areas are shown, A8b (A, B) and A8aD (E, F). The two prefrontal areas are displayed alongside posterior PGM (C, D). A8b receives projections from within or near several zones of the candidate default network including TPO and regions along the posterior cingulate (A30, A29a-c, A23V, and ProsSt). A8b also receives projections from parahippocampal area TH. A8aD displays overlap. PGM possesses some overlap, in particular for the injection in (C) that is in the ventral portion of PGM bordering A32V and A30. The position of the Ventral PGM injection site is shown by a red triangle in panel A to fully appreciate its location in relation to the borders of areas and in relation to the A8b injection pattern. The PGM injection falling fully within PGM reveals considerably fewer (if any) projections from ventral posterior cingulate and generally spares temporal association and parahippocampal cortex.

What is also revealed is that the present default-network-like candidate comprises zones beyond those typically hypothesized from the human neuroimaging literature including a single or multiple zones within and along the border of A47O/A13L/A13M. Projections from this cluster of areas are observed for both A10 (Figure 3A-C), TE3 (Figure 4E) and TPO (Figure 4A-B) injections. The candidate homologue of this zone is typically not well sampled in human fMRI studies because of susceptibility artifact and therefore would not have had the opportunity to nominate itself as a consistent component of the default network (but see Shulman et al., 1997).

Taken together these additional results illustrate that the apex transmodal network is more complex than the hypothesized five zones and further that the caudal to rostral gradient, while capturing general features of the organization, is not a complete description of the complex topology of the network which involves multiple interdigitated areas akin to the human (Braga and Buckner, 2017).

A question that may arise is whether the patterns revealed by the targeted analyses above capture unbiased properties given the several assumptions. To bolster confidence, as a final analysis all of the 143 available injections were subjected to a data-driven factor analysis. Consistent with the collective results, the second major factor broadly distinguished the apex transmodal network from sensory-motor hierarchies (Supplementary Figure 2). The one place where the distinction was not present is the region within and around auditory cortex that, as illustrated in Figure 4, has injections that display a mixed pattern. Of interest, when the injections within and near to auditory cortex are analyzed by themselves, the results reinforce separation of a preferentially auditory zone from a rostral temporal zone that participates in the transmodal network (Supplementary Figure 2 inset).

## Discussion

A recurring theme in comparative analysis of the marmoset brain as contrasted with larger New and Old World monkeys and human is that numerous areas, including temporal and prefrontal association areas, are conserved in kind and general relative positions to one another but often with less differentiation (Brodmann, 1909; Peden & Von Bonin, 1947; Burman et al., 2006; Burman & Rosa, 2009). The present analyses of marmoset tracer injection data reveal, at the broadest level, that the major distributed networks that have been postulated in the human are likely conserved with the same basic organizational motif and relative positions to one another on the cortical surface. Many differences in detail are expected including specializations, expansions, and differences in the propensity of afferent projections to higher-order association areas, but that the general macroscale organization of networks may be conserved between the marmoset and human is important for conceptual and practical reasons.

### A Macroscale Gradient of Networks is Present in the Marmoset

Analysis of large numbers of widely distributed tracer injections reveal a gradient of anatomical networks involving caudal to rostral frontal areas (Figure 2). While all of the networks conform to a basic motif of anterior and posterior cortical areas, there are notable transitions between networks that might provide insight into function and how simpler evolutionarily old motifs become expanded into the large-scale distributed networks that underpin higher cognition in primates.

In particular, the two most caudal networks (motor: A4ab-A3a and caudal premotor: A6DC) are more locally organized than the remaining networks. Versions of this transition have been noted previously in the marmoset (e.g., Bakola et al., 2015) and the macaque (Caminiti et al., 2015). In a thorough discussion of the fronto-parietal network archetype, Averbeck, Caminiti and colleagues explore extensive macaque anatomical data to describe separable fronto-parietal networks that have varying degrees of parallelism (Averbeck et al., 2009; Caminiti et al., 2015). They observed progressively more distributed networks as the fronto-parietal hierarchy was ascended with a network involving macaque PGm interconnected with a LIP/VIP/Opt cluster and a set of diverse prefrontal areas (including 8a, 8b, 45a/b, and 46v). The present transition from the caudal premotor network to the rostral premotor network, in Figure 2, may be similar to the transition that Caminiti and colleagues describe in the macaque (see also Margulies and Petrides, 2013).

Further convergent analysis comes from macaque functional connectivity data. Margulies et al. (2009), in a comprehensive analysis of the posterior midline in macaque, noted a transition from a sensorimotor-related network to a more extensive network linked to cognitive function. Recent analyses in the marmoset using functional connectivity also reveal support for transitions from a localized somatomotor network, to a more distributed frontoparietal network candidate of the macaque FEF-MT network, and then to more rostral prefrontal networks (Ghahremani et al., 2017).

At the apex of the progression observed here in the marmoset was a transmodal network that involved frontopolar A10 and rostrotemporal TE3. The network was considerably more extensive than revealed by the initial anterior-posterior target areas, with the network likely including extended components of rostral prefrontal (e.g., A11, Figure 3) and temporal association (e.g., TPO, Figure 4) cortex, as well as areas of dorsolateral (e.g., A8b and A8aD, Figure 5) and ventrolateral (e.g., see gray arrows in Figures 3A and 4E) prefrontal cortex. What was striking about this network is that it was converged upon by tracer injections distributed across multiple zones of cortex that mapped closely to predicted components of a homologue to the human default network.

### A Default Network Candidate in the Marmoset

The apex transmodal network involved the components expected of a homologue to default network - A in the human including: (I) frontopolar A10, (II) posterior midline A29a-c, A23, rostral A30, (III) rostral temporal association cortex TE3 / TPO / PGa/IPa, (IV) posterior parietal cortex Opt / PG, and (IV) parahippocampal cortex TH/ TF. Of these zones, the most ambiguous was Opt / PG.

Parietal association cortex received complex and diverse projections across all of the networks beginning with the rostral premotor network A6DR-PGM. In the human, the inferior parietal lobule and immediately adjacent regions contain a diverse set of juxtaposed networks that interact with distinct distributed networks involving visual and limbic areas (Vincent et al., 2008; Nelson et al., 2010; Yeo et al., 2011; Braga & Buckner, 2017; see also Culham & Kanwisher, 2001). In the macaque, injections to parahippocampal cortex are among the most informative for disambiguating the critical portion of Opt (e.g., Blatt et al., 2003; Lavenex et al., 2002; see Buckner et al., 2008 for discussion). Such injections are not yet available in the marmoset. Nonetheless, details of the presently available injections provide insight.

Frontopolar cortex A10 received minimal (but some) direct projections from posterior parietal cortex (similar to macaque). Adjacent A9 and A11 receive modest to strong projections (Figure 3E-F). Combined with the observation that the injections to rostral TE3 and caudal TPO label Opt/PG/LIP, these patterns converge to suggest that the A10-TE3 network includes a zone of posterior parietal cortex either through weak direct projections or, more likely, polysynaptic projections via its larger medial / orbital frontal network and distributed network with TE3 / TPO. Case CJ181-CTBr, involving A11, captured the pattern predicted of a default network candidate most closely and quite robustly.

Another feature of the default network candidate’s projection patterns is that they suggest separation from early to midlevel sensory processing areas, but the picture is not as clear as might be expected from the human default network. Supplementary Figure 1 illustrates a near complete dissociation between the default-network-like candidate and the A8aV-MT sensory-motor network, paralleling observations in the human (e.g., Fox et al., 2005; Fransson et al., 2005). But there were other examples of possible connections to sensory networks. For example, multiple cases showed some label in FST and MST (see Burman et al., 2011 for discussion). The projections were typically in the context of a broader region sending projections to A9/A10/A11 that involved rostral temporal association cortex through to the temporal pole. Possible other connections to higher-order visual areas have been noted previously and include projections from ProSt (Burman et al., 2011).

One particularly interesting zone was the border of auditory cortex and the polysensory temporal area TPO. TPO injections recapitulate the complete (Figure 4A) or nearly complete (Figure 4B) candidate default network pattern. In addition, TPO injections label projections from auditory areas. It is not possible with the present data to definitively interpret this juxtaposition of a default-network-like pattern and simultaneously strong projections from sensory areas. There may be distinct zones of TPO or its border may be misestimated in ways that complicate interpretation of the injections given the tight anatomical transitions (Supplementary Figure 2). The border between TPO and adjacent areas in the Paxinos et al (2012) atlas has been considered for revision (Majka et al., 2018 Figure 10) and the present gradient of temporal lobe projections follows a path that progresses through and divides the TPO / TE3 region. Another possibility is that the marmoset’s default network - A homologue does not have clear differentiation from auditory sensory processing areas. While less differentiation is a possibility, given the balance of evidence, we still consider the network as largely transmodal. What is most interesting about this zone is the opportunity for experimental dissection.

Common marmosets are vocal primates that live in groups and co-parent providing the opportunity to study rich social behaviors (Miller et al., 2016) including productive and receptive (auditory) communication (e.g., Wang, 2013; 2018; Song et al., 2016; Eliades & Miller, 2017). That so much is known about the marmoset auditory system and paradigms have been developed to task the system with simple sensory stimuli through to more elaborate social stimuli (Wang, 2018; Nummela et al., 2017) is fortuitous given that the areas involved in auditory processing are closely juxtaposed to those that transition to transmodal default network candidates (Figure 4F; Supplementary Figure 2). In future studies, the same experimental window (e.g., Nishimura et al., 2018) may provide the ability to physiologically measure both auditory processing areas as well as components of the marmoset default network.

### Limitations

There are a number of limitations to the approach taken here. First, in relation to the goal of drawing homologies with the human default network, the present work is based on anatomical connectivity (with reference to architectonic and positional correspondence between species). The human default network was originally defined by task suppression in neuroimaging studies and later explored in detail using functional connectivity (e.g., Buckner et al., 2008; Raichle et al., 2015). The present analyses do not draw from physiological task suppression effects, which constitute the basis of complementary explorations (e.g., Liu et al., submitted).

Another limitation is that there were relatively few tracer injections available in several key zones to fully appreciate the organization of the apex transmodal network. Specifically, injections along the ventral posterior midline and parahippocampal cortex were absent from analysis. Only a single injection was available for rostral TE3, none for nearby temporal pole areas TE1 and TE2, and there was no posterior partner injection for A6DC (Figure 2). There were also no injections within the unexpected projection zone along A47O/A13L/A13M to further explore this region. Greater coverage of the cortex will be critical to resolve finer details of the default network candidate. The present results show candidate homology with default network - A. An open question is whether there is further organizational subdivision as recently observed in the human (Braga & Buckner, 2017) or whether the marmoset network is less differentiated. Given the complexity of the network (Figure 5) and a proposal for a default network - B candidate (Liu et al., submitted), these details will be revealing.

A final limitation, perhaps to be considered a general limitation of the field, is that the areal definitions were not always predictive of the projection patterns, and in some instances, to the degree relied on as units for quantification, may prevent identifying patterns. For example, the rostral portion of TE3 received dense projections from multiple areas of the A10 network, while the caudal portion of the area was sparred. A reasonable hypothesis is that TE3 is heterogeneous with the caudal portion more aligned to a sensory pathway and the rostral portion a transmodal zone with distinct projections (see Figure 4F). As another example, the posterior parietal zone encompassing LIP/PG/Opt was complex with projections often on the border of the three areas. There were gradients – the A47L-AIP network projections tended to be rostral to the A8aV-MT network projections, but there was no simple alignment of projection gradients and areal boundaries, possibly reflecting a complex relationship between connectivity networks and evolutionarily new sectors of cortex (see Rosa & Tweedale, 2005 for discussion). Both PG and TPO in the marmoset are among the most functionally coupled areas to diverse cortical fields (Ghahremani et al., 2017) and within or near evolutionarily expanded cortical regions (Chaplin et al., 2013).

In the future large sets of injections fully mapped to the cortical mantle may provide a means to test alternative models of network organization – models that anchor from areal definitions and boundaries, and models that examine the projection patterns in relation to gradients and interdigitated patches. The availability of large databases of mapped projections and expression patterns, in common coordinate systems and annotated by traditional architectonic areal definitions, should allow the data to reveal its organizational features. Several efforts underway are working towards such opportunities (e.g., Okano et al., 2016; Majka et al., 2016; Liu et al., 2018).

## Conclusions

Anatomical analyses of the networks that progress along a caudal to rostral gradient in the common marmoset reveal macroscale homology with the human. Just as the relative positions of primary sensory and motor areas are conserved (Krubitzer, 2007), it appears that the broad ordering and relations of regions within distributed association networks may also be conserved. This finding has two distinct, important implications. First, the candidate homologies provide an opportunity to study how association networks form and differentiate in ways that may go awry in neuropsychiatric and developmental stress models. That the networks share similar features to the human provide specific opportunities to explore translatable properties of higher-order association networks. Second, the homologies provide a window into the organization of association networks that evolved at least 40 millions years ago, including a candidate homologue of the default network — perhaps best to be referred to as the apex transmodal network to minimize assumptions about its functional domain. An intriguing, but unproven idea, is that the apex transmodal network may have expanded and specialized for advanced human cognitive and social capabilities. The documented homologies provide an avenue to better understand how circuits, similar to those studied extensively in the human, function and interact mechanistically using modern physiological and molecular-genetic approaches.

## Methods

The present methods are based on the publically available data provided by the Marmoset Brain Architecture Project (http://www.marmosetbrain.org). This tremendously valuable open resource provides a comprehensive database with 143 retrograde tracer injections in the common marmoset (as of 8/2018) including published and unpublished injections all visualized on common flat map and surface volume projections, and annotated based on the Paxinos et al. (2012) atlas. Relevant methods are briefly described. The reader is referred to the original description in Majka et al. (2016) for details of how the data were digitized and brought into a common atlas framework. Figure 1 illustrates the visualization format used here.

For the previously published cases, the original reports describe the projection patterns in detail (e.g., Burman et al., 2011; 2014a; 2014b; Reser et al., 2013). Here we contrast between putative separate networks and between spatially distant injections in anterior and posterior cortical zones, as described below. When injection details are described, including if an injection invaded an adjacent area, that description was derived from the annotation of the injection in the Marmoset Brain Architecture Project archive (http://www.marmosetbrain.org).

### Subjects

All tracer injections were from the common marmoset (*Callithrix jacchus*). Four main types of fluorescent retrograde tracers were employed as described in Majka et al. (2016): flouroruby, FR; flouroemerald, FE; fast blue, FB; and diamidino yellow, DY. Several cases were not included in that original report and have subsequently been processed and uploaded to the online open resource. These cases additionally utilized tracers: choleratoxin subunit b green, CTBgr, and red, CTBr (for example, Cases CJ-CTBgr, CJ181-CTBr, and CJ180-CTBr). For the present analyses, 31 injection cases are the primary materials. Of these 12 are documented in prior publications (Burman et al., 2011; 2014a; 2014b; Reser et al., 2013) and the remainder pulled from the online database (Majka et al., 2016; http://www.marmosetbrain.org). In some instances, the maps are flipped from their original report to appear consistently in the space of the right hemisphere (e.g., Case CJ110-FE in the open release is flipped from its original presentation as Case 5 in Burman et al., 2014b).

### Analyses of the gradient of networks

In order to identify candidate networks organized throughout marmoset cortex, we anchored from the hypothesis that there might be sequential networks that appear one after the other progressing from primary cortices through to higher-order transmodal association cortex (Margulies et al., 2016). Given many details of network organization are not known, especially for the distributed networks that populate rostral portions of frontal cortex, this endeavor must be an approximation. Networks were sought that have separable long-distance connections (e.g., to posterior cortex). Less emphasis was placed on features of local connectivity that, for example in medial and orbital frontal cortex, can include complex projections between nearby areas forming partially segregated networks (Carmichael & Price, 1996; Öngür & Price, 2000; see also Burman et al., 2011). The emphasis here is on broad macroscale organization.

To first identify possible sequential networks, frontal injection sites were selected progressing through caudal to rostral areas: (1) primary motor cortex (M1), (2) caudal premotor area A6DC, (3) rostral premotor area A6DR, (4) one of the multiple candidates for the frontal eye field (FEF) area A8aV, (5) prefrontal area A47L, and (6) frontopolar area A10. The cascade along frontal cortex parallels the recent analyses in the human using functional neuroimaging methods (Braga and Buckner, 2017) but here operationalized in the marmoset based on anatomical tracer injections.

#### Assumptions behind selection of frontal injections

There were several assumptions in selecting this sequence. First, A4ab (primary motor cortex, M1) was assumed to be the lowest level. Second, A10 within the frontal pole was assumed to be at the highest transmodal level. Third, caudal premotor area A6DC was assumed to be at a relatively lower level than rostral A6DR (which directly followed M1). This relative positioning of A6DC and A6DR stems from the detailed analysis of the motor system in Bakola et al. (2015) where A6DC was demonstrated to have preferentially stronger projections to M1 relative to A6DR, and A6DR relatively stronger projections to posterior medial cortex (e.g., A23a-c), prefrontal, and PGM (see detailed analysis in Burman et al., 2014a; 2014b). Recognizing that there is debate about the exact homologue(s) of FEF, A8aV was selected because of its distinct network connectivity (Burman et al., 2006) and cursory parallels with network organization in the macaque (Maunsell & Van Essen, 1983; Ungerleider & Desimone, 1986) and human (Corbetta & Shulman, 2002; Fox et al., 2006). The placement of A8aV in its specific position in the sequence is not firm; while it is assumed to come after A6DC, it is possible that it is parallel or orthogonal to A6DR. Prefrontal A47L was situated in the sequence above all of the other areas, one level below A10. A47L was selected because of availability of multiple relevant injections and because it was positioned caudal to A10 and rostral to the other selected frontal areas.

#### Replication

For each selected frontal injection an independent injection in another animal was identified that substantially replicated the distributed pattern of projections. The initial injection / replication pairs were as follows: A4ab (Cases CJ78-FB and CJ173-DY), A6DC (Cases CJ111-FR and CJ112-FR), A6DR (Cases CJ110-FE and CJ116-FR), A8aV (Cases CJ125-FR and CJ75-DY), A47L (Cases DJ73-FB and CJ181-DY), and A10 (Cases CJ70-DY and CJ178-CTBr).

#### Posterior injections

In order to bolster confidence that the frontal injections are capturing distinct networks, the networks were all corroborated with an independent injection in one of the main posterior input areas. A4ab (primary motor cortex) was corroborated with an injection of A3a (primary somatosensory cortex; Case CJ170-FB); A6DR with an injection to medial parietal area PGM (Case CJ80-DY); A8aV with an injection to extrastriate visual area MT (CasesCJ56-FB); A47L with an injection to parietal association area AIP (Case CJ173-CTBgr), and A10 with an injection to the rostral portion of temporal association area TE3 (Case CJ180-CTBr). There was no available injection to corroborate A6DC (the only available injection in the relevant area, PE, was on the border invading somatosensory area 1/2, e.g., Case CJ173-CTBr). For simplicity, the networks linked to each anterior-posterior injection pair is referred to by its initial targeted areas (e.g., A10-TE3 network) recognizing the full network is considerably more extensive (e.g., the A10-TE3 network likely includes A11 and TPO, among many other areas).

### In depth examination of the default-network-like candidate

As the results will reveal, the marmoset possesses a set of separate networks progressing from primary motor cortex through to frontopolar cortex. The apex transmodal network associated with A10 includes many of the regions expected of a human default network homologue. To explore this network further, five additional injections were examined in and adjacent to A10. In human, the large region of medial prefrontal cortex that is involved in the default network includes area 10 (human 10m, 10r, and 10p), area 9, as well as anterior cingulate areas human 24/32ac (based on Öngür et al., 2003 as analyzed in Buckner et al., 2008). The additional injections allowed the areas in the vicinity of A10 to be explored more thoroughly. These injections included: caudal A10 (Case CJ178-CTBgr; with possible slight invasion of A46); medial frontopolar A10 (Case CJ73-FR), A32V (also known as 10mc, Burman & Rosa, 2009); A11 (Case CJ181-CTBr with some leakage up the track in dorsolateral prefrontal cortex), and A9 (Case CJ70-FB). As the investigation progressed, additional tracer injections were examined to resolve and expand on relevant patterns.

## Acknowledgments

The Marmoset Brain Architecture Project (http://www.marmosetbrain.org) was the critical open resource that provided the tracer injection data. We thank Marcello Rosa and CiRong Liu for valuable discussion. Katie Insel commented on an early draft and Hannah Becker provided assistance with the references. This work was supported by NIH grant P50MH106435.

## Supplementary Materials

**Supplementary Figure 1.**
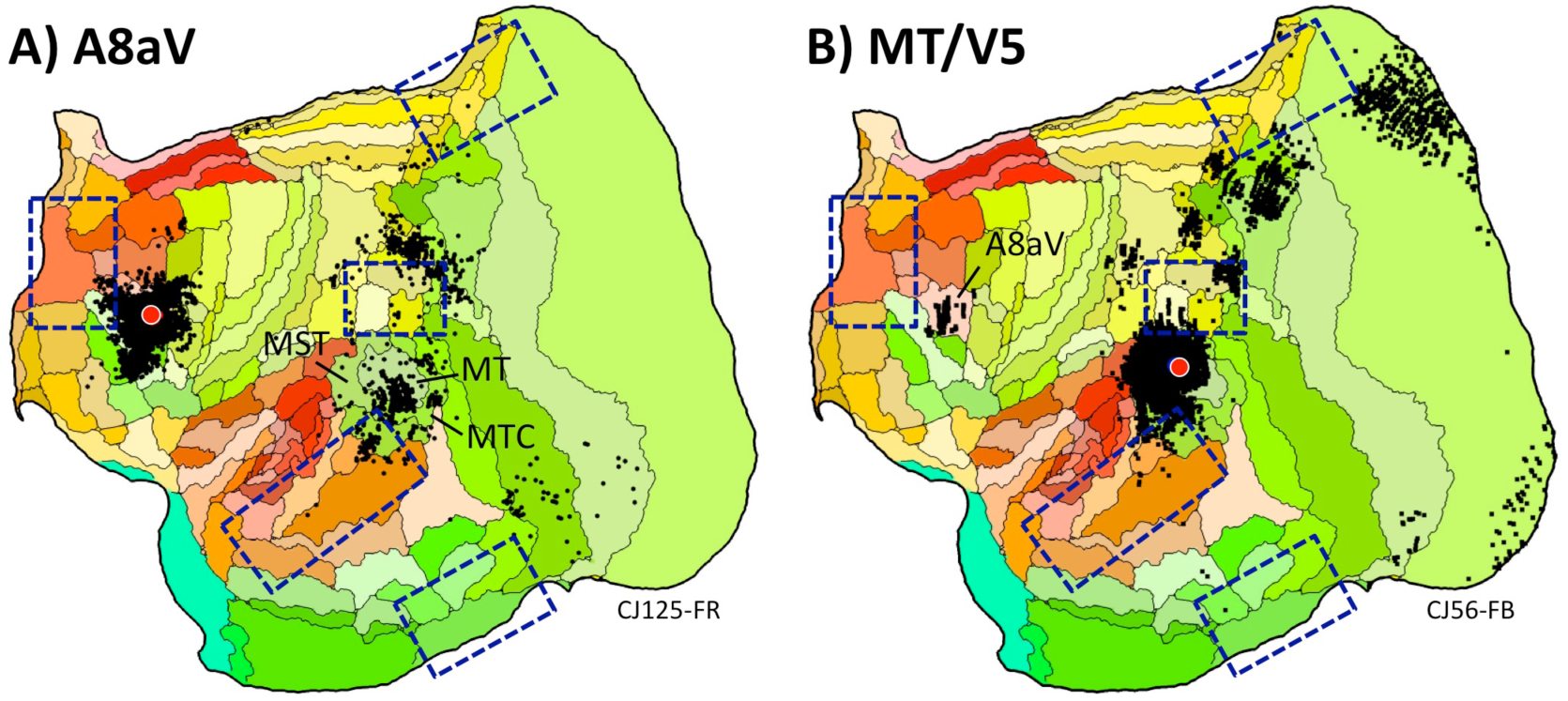
Evidence for separation from a sensory-motor network. Displayed in (A) is an injection to a FEF candidate in the marmoset (A8aV) contrasted with an area MT injection in (B). The two maps illustrate that the anterior frontal and posterior areas are connected to each other. Critically, the patterns of connections are quite separate from the candidate homologue of the human default network – A (highlighted by blue rectangles).

**Supplementary Figure 2.**
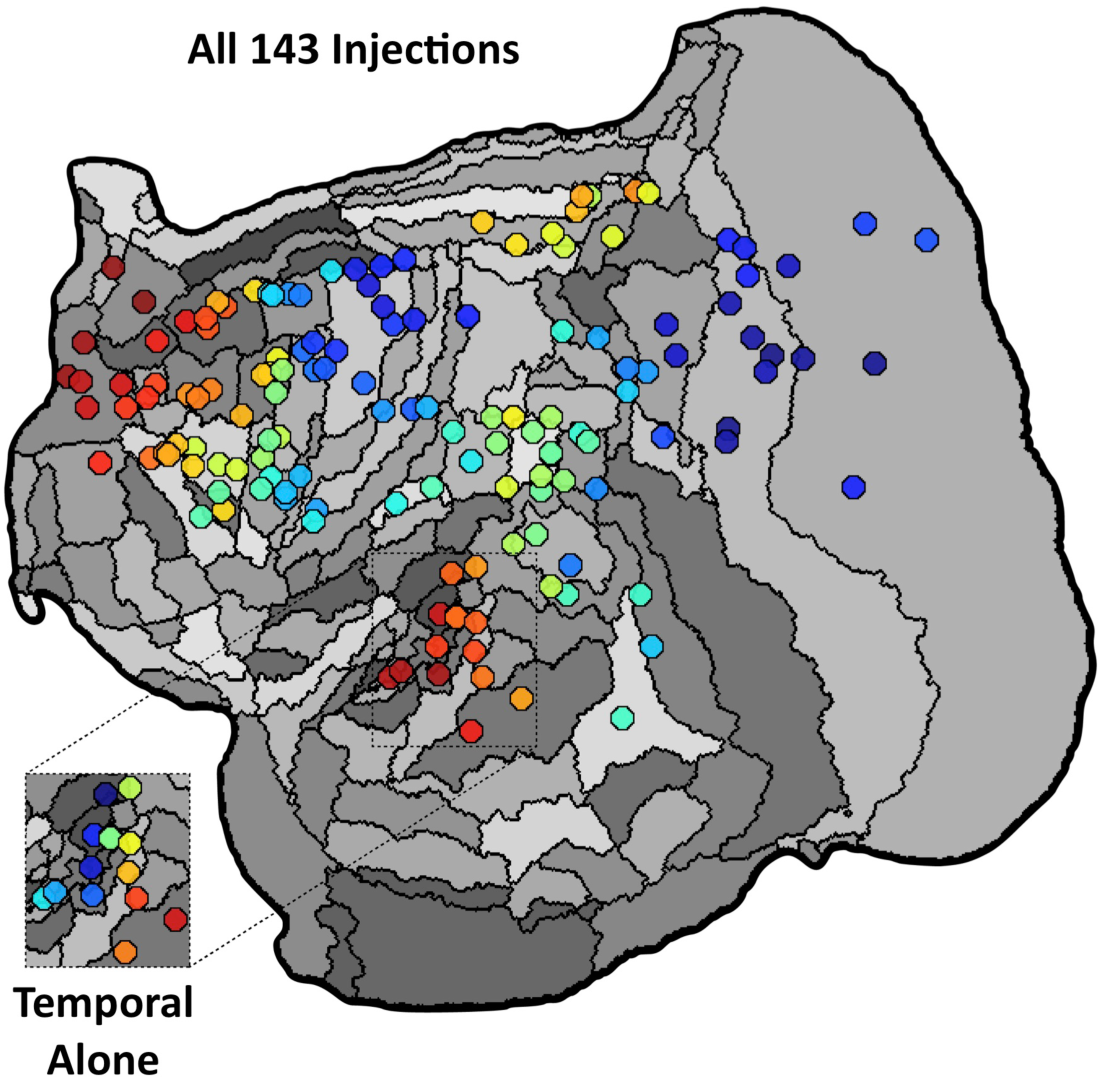
Transmodal association cortex is distinguished from sensory hierarchies when analyzed by unbiased factor analysis. Displayed are all 143 injection locations from the Marmoset Brain Architecture Project archive as of August, 2018 color coded based on their association with the major factor that distinguishes projection patterns in the marmoset cortex. The main figure presents the rank order of all injections, derived from diffusion map embedding of a similarity matrix calculated across flatmap connectivity patterns. While the first factor differentiated visual from somatosensory/motor areas, here we show the second factor corresponding with the spectrum from primary to apex areas. Injections displayed with red-yellow colors show the strongest weighting, while blue colors the opposite. The inset image presents a secondary analysis restricted to temporal lobe injections, in which the first factor demonstrated the gradient between auditory regions and areas TPO, PGa/IPa, and TE3. Code is available at [link to be made available upon publication].

